# Prioritizing Stability-enhancing Mutations using a Protein Language Model in conjunction with Physics-Based Predictions

**DOI:** 10.1101/2025.10.28.683791

**Authors:** Emily R. Rhodes, Guido Scarabelli, Jonathan Jou, Jacob Byerly, Kayla G. Sprenger, Eliud Oloo

## Abstract

Mutational protein engineering, as currently practiced in the biotechnology and pharmaceutical industries, is both tedious and expensive. Computationally driven protein design has the potential to expedite the process and generate high-quality variants at a lower cost. Using datasets of 174824 mutations for 180 proteins, we benchmark the effectiveness of a protein language model (PLM), Evolutionary Scale Modeling (ESM), alongside a physics-based method (PBM), namely molecular mechanics energies with generalized Born and surface area continuum solvation (MM/GBSA), as triaging tools for identifying and prioritizing target positions and specific mutations that improve protein thermodynamic stability. We found prediction biases in each method but also determined that these biases can be mitigated by applying the two methods in a complementary manner. We propose a hybrid mutation prioritization and selection strategy that achieves better accuracy than either method alone. Through re-ranking, the combined prioritization strategy attained a higher overall average ROC AUC of 0.744 across the dataset compared to either MM/GBSA alone (0.684) or ESM Log Odds alone (0.597).

## 1. Introduction

Protein stability assessment is a key feature for the discovery of new biological and pharmaceutical products, since it affects properties such as solubility, functional fidelity, expression yield, enzymatic activity, and shelf-life.^1–3^

Directed evolution is the traditional experimental approach used to generate more stable variants for proteins and has been successfully used in many enzyme design and protein engineering applications.^4,5^ However, the need for multiple rounds of genetic recombination and screening can require significant time and experimental costs in order to generate variants with the desired characteristics. Computational methods that reduce the cost and time required by such experimental approaches would thus be desirable.

### 1.1. Computational Protein Design Benefits and Obstacles

Most computational protein design methods are either knowledge-based, leveraging statistical correlation to predict experimental outcomes, or physics-based, approximating underlying physical forces directly. Some of these include tools such as evolutionary information and statistical potential,^6,7^ force fields,^8^ structural modeling,^9–11^ and machine learning (ML)^12–14^, specifically support vector machines.^15–18^ A common modern approach in protein engineering involves using ML models—either pre-trained^19^ to predict mutations, or foundation models fine-tuned for a specific engineering application such as antibody engineering.^20^ Both approaches depend heavily on the availability of large amounts of high-quality data that is sufficiently related to the intended use-cases; limited or low-quality data will reduce the predictive accuracy of both methods. In contrast, a physics-based model utilizes a force field to estimate the energy cost associated with a physicochemical change.^21^ By estimating known physical forces at the atomistic level, physics-based methods are designed to predict experimental observables for not only the structural and experimental dataset they were built upon, but also new, unseen structures and conformations. Limits in our descriptions of these physical forces and computational resources prohibit physics-based methods from readily scaling to large macromolecular systems.

In this work, we explored a complementary approach: pairing a foundation model with a physics-based model. This layered approach predicts specific properties without the need for retraining, leveraging the speed of ML and the generalizability of physics-based models for diverse use cases. Toward this end, we focused on two widely used computational methods: a protein language model (PLM) referred to as Evolutionary Scale Modeling (ESM) and a physics-based method (PBM), namely molecular mechanics combined with generalized Born and surface area continuum solvation (MM/GBSA).

### 1.2. Protein Language Models

PLMs are a form of machine learning which excels at predicting reasonable protein sequences and structures by learning amino acid contexts from large datasets.^19,22–25^ When applied to proteins, PLMs predict masked amino acids in sequences analogous to words in sentences. ESM-1b illustrates this concept by estimating amino acid probabilities at specific sequence positions.^23^ Trained on UniParc’s evolutionary data^26^ using self-supervised learning, ESM-1b discerns patterns in protein sequences. With 650 million parameters and trained on 250 million sequences (totaling 86 billion amino acids), this model predicts protein functions, and mutations by uncovering key biochemical properties of amino acids. ESM-1b has a parameter:data ratio of 2:5, where 100 million parameters were fit while training across 250 million sequences, limiting overfitting.

While ESM-1b is not designed or optimized to predict specific biophysical properties like stability, the information it provides on the probability of a mutation compared to wild type can be applied as a general indication of mutations that are evolutionarily plausible: likelier mutations must not only adequately retain existing function, but also impart some improvement in fitness over wild type.^20^ In general PLMs are able to recreate sequences, indicating that the model is trained to form an internal representation of the information contained in protein sequence data.

### 1.3. Physics-Based Models

PBMs take into account chemical forces to estimate a change in free energy. The state-of-the-art method, free energy perturbation (FEP), makes fewer assumptions and describes the physics in a more complete manner when compared to MM/GBSA which sacrifices some accuracy to make the calculation cheaper and faster. However, both methods attempt to estimate free energy in a similar manner.

Different states of the protein (folded versus unfolded, and in the absence or presence of incorporated mutations) are used to estimate the change in free energy associated with transforming one residue into another, as described in **Eqn. 1** below:

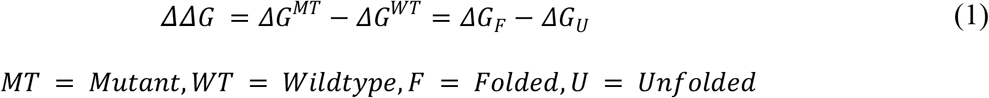

where *ΔΔG* represents the change in free energy of folding due to mutation, *ΔG*^*MT*^ is the free energy change associated with folding of the mutant, Δ*G*^*WT*^ is the free energy change associated with folding of the wildtype, *ΔG*_*F*_ is the free energy change associated with making a mutation in the folded protein, and *ΔG*_*U*_ is the free energy change associated with making a mutation in the unfolded protein. Experiments measure *ΔG*^*MT*^ and *ΔG*^*WT*^, while PBMs use *ΔG*_*F*_ and *ΔG*_*U*_. Although this process is non-physical, its estimates align with observed changes in free energy in physical systems (*R*^2^ = 0.94).^21,27^

### 1.4. Model Bias

Both MM/GBSA and ESM-1b have bias due to their construction. MM/GBSA employs an implicit solvent model, which might contribute to an overestimation of charged interactions on the protein surface and therefore overpredict the effect of mutations involving residues with charged side chains as seen in **Table 1**. It also does not account for major protein structural rearrangement which could occur as a result of mutation. ESM-1b does not have a clear representation of the three-dimensional structure and therefore could overpredict the effect of mutations involving residues with hydrophobic side chains because the surface and core of the protein have different local contexts, similarly observed in the results from **Table 1**.

**Table 1.** Capture of stabilizing mutations by MM/GBSA and ESM-1b by categories of amino acid mutations for the Mega Dataset.

|  | Exp. Hit | MM/GBSA |  |  | ESM Log Odds |  |  |
| --- | --- | --- | --- | --- | --- | --- | --- |
|  |  | Top 20% (Protein) | Top 5 (Position) | ROC AUC | Top 20% (Protein) | Top 5 (Position) | ROC AUC |
| Positive to Negative | 113 | 39.8% | 8.85% | 0.707 | 40.7% | 35.4% | 0.572 |
| Negative to Positive | 237 | 48.9% | 32.9% | 0.691 | 31.6% | 36.7% | 0.614 |
| Positive to Non-charged Polar | 248 | 47.6% | 16.1% | 0.597 | 23.8% | 22.6% | 0.556 |
| Negative to Non-charged Polar | 468 | 52.4% | 15.8% | 0.668 | 25.6% | 22.6% | 0.593 |
| Non-charged Polar to Positive | 238 | 53.4% | 36.6% | 0.686 | 60.5% | 37.0% | 0.644 |
| Non-charged Polar to Negative | 150 | 21.3% | 14.0% | 0.667 | 67.3% | 46.7% | 0.667 |
| Negative to Hydrophobic | 896 | 54.6% | 30.2% | 0.636 | 20.5% | 15.8% | 0.584 |
| Hydrophobic to Negative | 148 | 27.0% | 9.46% | 0.803 | 48.6% | 27.7% | 0.707 |
| Positive to Hydrophobic | 639 | 55.2% | 29.1% | 0.657 | 19.7% | 19.1% | 0.543 |
| Hydrophobic to Positive | 139 | 44.6% | 30.2% | 0.727 | 42.4% | 30.2% | 0.662 |
| Non-charged Polar to Hydrophobic | 1130 | 51.3% | 34.5% | 0.686 | 39.5% | 22.6% | 0.611 |
| Hydrophobic to Non-charged Polar | 346 | 26.9% | 16.8% | 0.721 | 41.9% | 18.8% | 0.697 |
| Non-charged Polar to Non-charged Polar | 406 | 35.5% | 20.0% | 0.631 | 45.6% | 28.1% | 0.629 |
| Hydrophobic to Hydrophobic | 973 | 37.4% | 29.4% | 0.694 | 41.8% | 25.0% | 0.654 |
| Negative to Negative | 44 | 40.9% | 9.09% | 0.621 | 88.6% | 56.8% | 0.804 |
| Positive to Positive | 31 | 80.6% | 16.1% | 0.664 | 54.8% | 22.6% | 0.482 |
| Any to Proline | 131 | 8.40% | 3.05% | 0.832 | 53.4% | 48.9% | 0.822 |
| Proline to Any | 224 | 29.9% | 22.3% | 0.637 | 48.7% | 19.2% | 0.691 |
| Any to Cysteine | 821 | 15.8% | 2.56% | 0.665 | 4.75% | 2.68% | 0.574 |
| Cysteine to Any | 1 | 0.00% | 0.00% | 0.324 | 100% | 0.00% | 0.757 |
| Small to Large | 927 | 49.6% | 25.6% | 0.673 | 28.3% | 16.4% | 0.605 |
| Large to Small | 861 | 20.3% | 5.69% | 0.636 | 26.8% | 23.2% | 0.510 |
| Any to Any | 7360 | 41.5% | 23.4% | 0.690 | 33.1% | 22.2% | 0.599 |

### 1.5. Complementary Features of PBMs and PLMs

While PBMs aim to capture well understood physical phenomena in a concise manner, there are features of physical systems not captured by these methods, namely charges are static preventing chemical reactions from being captured. On the other hand, PLMs learn from evolutionary data and therefore can learn nuance (ex: impact of charged mutations) from these systems that is not easily incorporated in a force field. Therefore, we posit that ESM-1b captures distinct biophysical or evolutionary information complementary to MM/GBSA, suggesting their combined application could improve mutation prioritization.

## 2. Methods

### 2.1. Benchmark Datasets

To assess ESM-1b and MM/GBSA in a stabilizing context, we used the Nisthal dataset and the Mega dataset. The first, the Nisthal Dataset,^28^ comprises data from a comprehensive automated site-directed mutagenesis and chemical denaturation study performed on the B1 immunoglobulin-binding domain of streptococcal protein G (Protein Data Bank (PDB) code 1PGA^29^), with measured thermodynamic stability properties. This dataset allowed us to evaluate ESM-1b’s ability to predict stabilizing mutations on a benchmark set of data consisting of changes in the free energy of folding for nearly every single residue variant for a small, 56-residue protein. These questions were then extended to a second, much larger dataset, the Mega Dataset,^30^ which contains data characterizing the impact of mutations on the stability of various proteins. From the Mega dataset, we selected proteins previously deposited in the PDB with more than 500 experimental stability measurements for a total of 179 proteins and 174010 benchmark stabilities. Within the experimental datasets, a stabilizing mutation was defined as one that leads to a decrease in free energy of more than 0.5 kcal/mol, while a destabilizing mutation was defined as one that leads to an increase in free energy of more than 0.5 kcal/mol.^31^

### 2.2. Evolutionary Scale Modeling - 1b Variant Prediction

To generate log-odds ratios from ESM-1b, we modified the prediction module of ESM-1b to return log-odds instead of simply returning the likeliest AA. ESM-1b was otherwise unchanged, and no model parameters were modified. A zero-shot prediction was attempted to understand how this model can perform “out of the box”.

### 2.3. Protein Preparation

The Mega Dataset includes AlphaFold-predicted structures of all proteins used in the experiments, which we used as initial structures for MM/GBSA simulations. The structures were prepared using Schrödinger’s Protein Preparation Workflow,^32^ which incorporates PROPKA^33,34^ for predicting pKa values and protonation states. PROPKA was run at a pH of 7.0 ± 2.0. Clusters of atoms within the hydrogen bonding network were optimized to achieve the lowest energy state, allowing for flips or stacking of the side chains. Finally, the structures, parameterized using the OPLS4^35^ force field, underwent energy minimization.

### 2.4. MM/GBSA

To estimate changes in protein free energy of folding, MM/GBSA calculations were performed using Schrödinger’s BioLuminate platform,^32^ which employs an implicit solvation model and the OPLS4^35^ force field. This approach balances speed and accuracy in estimating the energy required to mutate a given residue in a protein structure.^36^ The Prime energy function is used to calculate the change in stability of the protein due to the mutation with an implicit solvent term.

### 2.5. Data Post-Processing

The log-odds ratios from ESM-1b and the estimated change in free energy of folding from MM/GBSA were used to rank the mutations from best to worst. The likeliest mutation by log odds was interpreted to be the most favorable.^20^ When rankings are described for each methodology, the largest, positive log-odds ratio was given the top ranking while the largest, negative log-odds ratio was given the bottom ranking for ESM-1b and the largest, negative *ΔΔG* value was given the top ranking while the largest, positive *ΔΔG* value was given the bottom ranking for MM/GBSA. The rankings were converted to percentiles where the best mutations had low percentile values and vice versa.

Exclusively for truth table analysis, the computationally predicted “beneficial” mutations were defined as the top 5% of mutations ranked using either MM/GBSA, or for ESM-1b. Conversely, the computationally predicted “detrimental” mutations were defined as the bottom 5% of mutations ranked using either MM/GBSA, or for ESM-1b. These classifications were guided by insights sensitivity, specificity and accuracy analysis from the Nisthal Dataset (**Fig. S1**).

### 2.6. Combining ESM-1b and MM/GBSA

The percentiles from ESM-1b and MM/GBSA we combined using linear regression. This is described in **Eqn. 2** below:

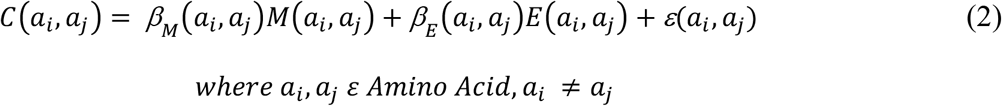

The coefficients *β* _*M*_ (*a*_*i*_*j a*_*j*_) and *β*_*E*_ (*a*_*i*_,*a*_*j*_) as well as the constant *ε* (*a*_*i*,_ *a*_*j*_) were calculated using Ordinary Least Squares (OLS), fitting to the experimental percentile distribution. *M*(*a*_*i*_,*a*_*j*_) is the percentile from the MM/GBSA calculation that represents mutating from i (amino acid) to j (amino acid). *E*(*a*_*i*_,*a*_*j*_) is the percentile from the ESM log odds calculation that represents mutating from i (amino acid) to j (amino acid). *C*(*a*_*i*_,*a*_*j*_) is the combined percentile calculated using the parameters described.

## 3. Results

### 3.1. Experimental dataset coverage and agreement

Due to experimental limitations, the Nisthal Dataset excludes mutations from tryptophan in the wildtype, as well as mutations to cysteine or tryptophan. (**Fig. 1A**). Furthermore, the wild-type sequence of streptococcal protein G lacks arginine, histidine, serine, cysteine, and proline residues, and mutations from these amino acids are also absent from the dataset. In contrast, the Mega Dataset offers near-complete coverage of all mutations, excluding occasional unobserved mutations (**Fig. 1B**). To compare positions on equal footing, only positions with complete experimental data for all mutations were included in the analysis (**Fig. S2**).

**Figure 1.**
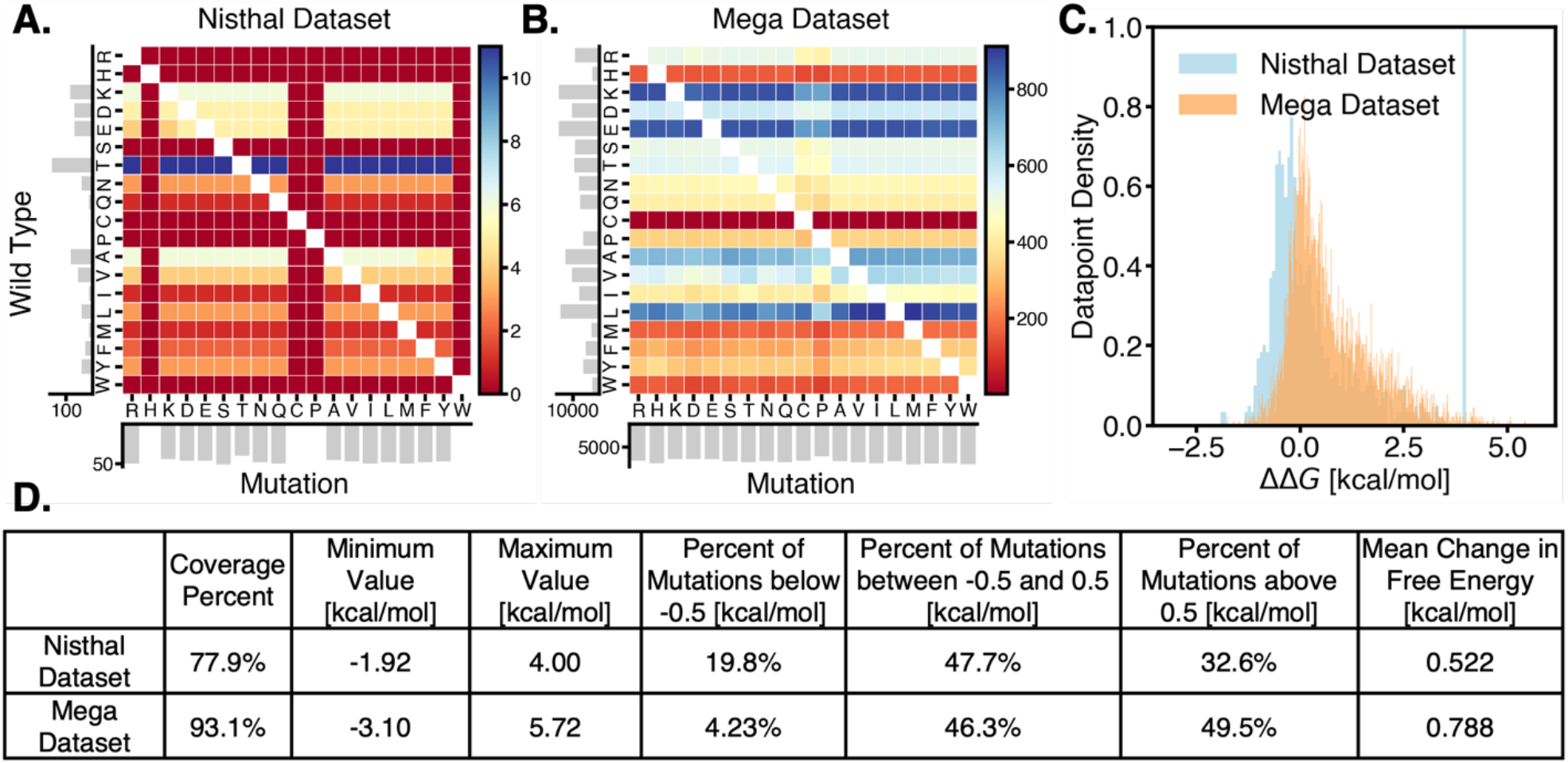
Experimental dataset coverage and values. A) Nisthal Dataset mutation coverage of the B1 immunoglobulin-binding domain of streptococcal protein G (PDB code 1PGA), with colors indicating mutation counts and gray bars indicating the sum of mutations in each respective column/row. B) Mega Dataset mutation coverage of all the proteins studied in this dataset, with colors indicating mutation counts and gray bars indicating the sum of mutations in each respective column/row. C) Changes in free energy, ΔΔG, represented between the Nisthal Dataset and Mega Dataset for all mutants. D) Experimental dataset coverage and population values categorized by ΔΔG.

When comparing the two datasets, we observed an overlap in measured free-energy changes. However, we also saw a larger fraction of mutations assayed as stabilizing (and a smaller fraction of mutations assayed as destabilizing) in the Nisthal Dataset as compared to the Mega Dataset, despite both measuring the same property (*ΔΔG*) (**Fig. 1C-D**).

Overall, the Nisthal Dataset has 77.9% coverage of mutations to the B1 immunoglobulin-binding domain of streptococcal protein G, and the Mega Dataset has 93.1% coverage of mutations to all proteins within the selected experimental data subset. Both datasets have a similar mean change in free energy. While the Mega Dataset has a wider range of changes in free energy (both a higher maximum value and a lower minimum value), it also includes more proteins and a higher coverage of all the mutations. These two datasets are used in conjunction to evaluate each method in a variety of contexts and against two different experimental methods for measuring stability.

### 3.2. Individual method performance: identifying stabilizing mutations and *mutable sites*

First, assessing the methods individually, we evaluated the degree of separation between the populations of stabilizing and destabilizing mutations using each computational method. As anticipated, MM/GBSA (**Fig. 2A, Fig. S3A**) separated the populations more distinctly compared to ESM-1b (**Fig. 2B, Fig. S3B**). While both methods significantly distinguished the stabilizing from destabilizing mutations, as indicated by low KS test p-values, in both datasets MM/GBSA showed greater separation, evidenced by larger KS test statistic, where the MM/GBSA KS test statistic was 0.454 for the Mega Dataset and 0.709 for the Nisthal Dataset and the ESM Log Odds KS test statistic was 0.229 for the Mega Dataset and 0.255 for the Nisthal Dataset (**Fig. 2C, Fig. S3C**).

**Figure 2.**
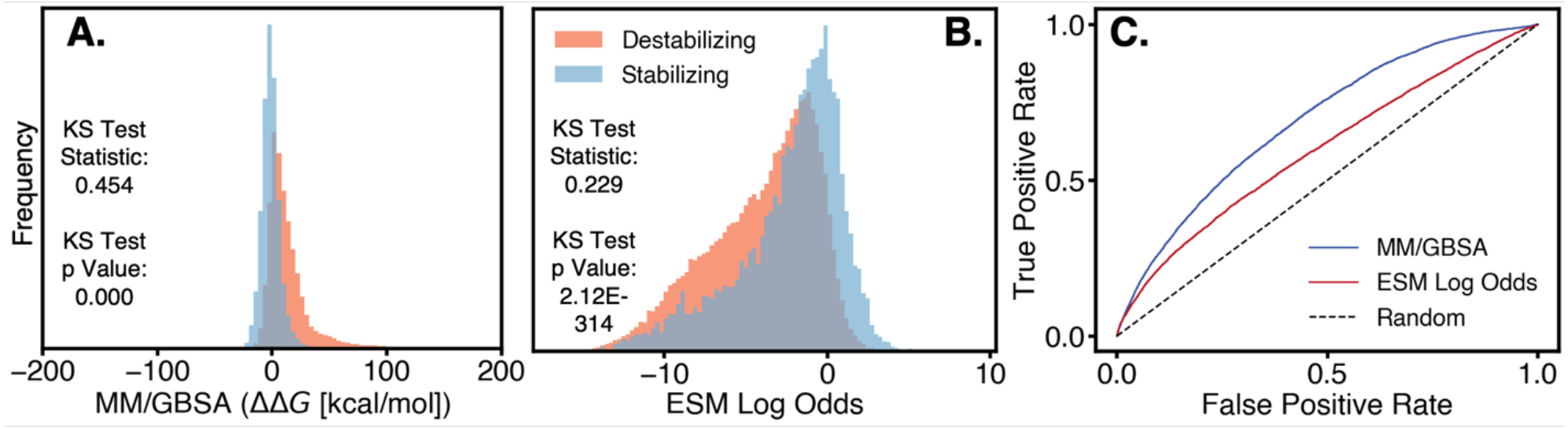
Distinguishing stabilizing mutations. The population of mutation values measured by A) MM/GBSA, and B) ESM log-odds ratios for stabilizing (blue) and destabilizing (orange) mutations, across the entire Mega Dataset. Significant differences in populations are indicated by the p-value of a KS test and the degree of separation by the KS statistic. C) Receiver operating characteristic curve for both MM/GBSA and ESM log-odds ratios.

In the case of ESM-1b, stabilizing mutations yielded slightly higher log-odds ratios, on average, than destabilizing mutations. Though the difference was small, the direction of this shift matched expectations. This trend was observed both across the entire Mega Dataset (**Fig. 2B**) and in the Nisthal Dataset (**Fig. S3B**). Interestingly, the distributions of stabilizing mutations for both datasets showed a tail skewing toward more negative log-odds ratios. This tail supports our hypothesis that mutations may be disfavored not because of lower stability, but instead mere evolutionary undersampling. Furthermore, evolution selects for fitness, a measure which encompasses not only fold stability, but also binding affinity, interdomain flexibility, and binding selectivity, among others.

This trend was reinforced by assessing both the linear (Pearson correlation) and monotonic (Spearman correlation) relationships between the computational and experimental mutation rankings (**Fig. S4A**). MM/GBSA consistently outperformed ESM-1b (**Fig. S4A-D**). When ranking mutations at each position, all methods performed less effectively compared to ranking mutations across the entire protein (**Fig. S4A, Fig. S5A-B**). The truth table further illustrates ESM-1b’s limitations. Overall, MM/GBSA demonstrated a higher specificity, sensitivity and accuracy than ESM-1b across both the Mega Dataset (**Table S1**) and Nisthal Dataset (**Table S2**).

In an effort to identify *mutable sites*, we hypothesized that positions with low standard deviation—indicating a narrow range of computed computational values—would qualify as suitable *mutable sites* to target for design of stability improving substitutions, as they can likely accommodate a greater variety of mutations with minimal disruption. Stability hotspots were classified using the experimental data as sites where stabilizing mutations are frequently favored, as defined by majority of mutations meeting the ΔΔG threshold values and were thus labeled as stabilizing positions. Conversely, destabilizing positions were categorized as those where most mutations fell into the “destabilizing mutation” category as defined by experimental ΔΔG values. MM/GBSA more effectively separated the two populations (stabilizing and destabilizing positions) than ESM-1b across both the Mega Dataset (**Fig. S6**) and Nisthal Dataset (**Fig. S7**).

### 3.3. Individual method performance: mutation categories

To analyze prediction performance based on the type of mutational change, we categorized mutations by the physical and chemical properties of the wildtype and mutant residues involved (**Table 1**). We then employed ESM-1b and MM/GBSA to rank the mutations according to the computational values (mutation-induced free energy changes for MM/GBSA, and ESM log-odds ratio for ESM-1b). We then determined the percent of experimentally verified stabilizing mutations captured in both the top 20% of mutations (across the whole protein) and top 5 mutations (per position) for each category by each computational prediction method. Simultaneously we plotted the receiver operating characteristic (ROC) curve and reported the area under the curve (AUC) for each method, providing insight into which method performs best at prioritizing mutational substitutions of a given type. For each metric, applied on the Mega Dataset, MM/GBSA performed the best overall which is evidenced by the results in the ‘Any to Any’ row of **Table 1**.

When looking at the number of experimental hits in each category, one of the top categories is “Non-charged Polar to Hydrophobic” mutations with 1130 experimental hits out of the total 7360. Over 50% of these hits are captured by MM/GBSA in the top 20% of mutations. Several categories with a large number of hits are favored by MM/GBSA which may be a reason why MM/GBSA performs better, as a whole, compared to ESM Log Odds; MM/GBSA inflates the ranking of the mutations in the categories where experimental hits tend to be more likely. If there was truly no bias and each method performed equally well across all groups, we would expect to see roughly the same percentages in the “Top 20% (Protein)” column across the groups.

Using ROC AUC as the top metric for success, MM/GBSA performs better in each individual category except “Negative to Negative”, “Proline to Any”, and “Cysteine to Any” mutations. However, as indicated by the number of experimental hits and the “Top 20% (Protein)” and “Top 5 (Position)” columns this does not describe the complete picture. For the “Cysteine to Any” group, there is only one experimental hit and therefore there is no conclusive evidence to suggest one method outperforms the other. For the “Negative to Negative” group, as well as other “…to Negative” groups, ESM Log Odds indicates superior performance in the “Top 20% (Protein)” and “Top 5 (Position)” columns as well (compared to other groups), suggesting that ESM Log Odds also has some bias towards these mutation types. Lastly, the “Proline to Any” category performance suggests that these groups can be broken down into amino acid identity types to reveal other mutation types with enhanced performance using ESM Log Odds.

Looking at the “Top 20% (Protein)” and “Top 5 (Position)” columns, while ESM-1b performed worse at identifying mutations to hydrophobic amino acids compared to MM/GBSA, it typically identified mutations to charged amino acids with equal or better success rates than MM/GBSA, suggesting an area of weakness for MM/GBSA and possibly an opportunity for complementary application of both approaches. Furthermore, MM/GBSA comparatively struggled to capture mutations to proline or from relatively large-sized to small-sized amino acids. When examining solely the Nisthal Dataset, MM/GBSA showed similar drawbacks (**Table S3**).

Overall, the collective information across the columns in this data suggests biases where each method is capturing more or less mutations by shifting the values for a given category relative to all the other values. Additionally, these biases can be illuminated by analysis of categories of mutations but also through individual amino acid identity as seen with the “Proline to Any” category. To harness the strength of each tool and use a complementary approach involving both methods, the biases must be taken into account.

Each row is a different category of mutation where the first word describes the wildtype residue which is then mutated to another amino acid described by the second word (for instance, “positive to hydrophobic” indicates the wildtype amino acid is positive and it is then mutated to a hydrophobic residue). The columns indicate the percent of hits the computational method captured in the “Top 20% (Protein”) and “Top 5 (Position)” as well as the “ROC AUC” for all mutations of a given category.

### 3.4. Computational method biases

Mutant ranks were converted to percentiles for normalized comparison across proteins. Percentile distributions relative to experimental stability data revealed method-specific biases towards certain mutation types. These biases provide an explanation for why methods perform well in particular categories, but do not translate that success into overall increased performance. By biasing a particular category of mutations as very probable, every other mutation must be subsequently less probable and therefore over-estimates the favorability of a certain category of mutations (MM/GBSA values or ESM log odds values), increasing the performance in this category at the cost of a drop in performance for every other category. As an example, the mutations from negative amino acids to negative amino acids are heavily biased by ESM-1b (**Fig. 3A**) explaining why this method captures these mutations more rapidly, however, it also orders these mutations more accurately than MM/GBSA as seen by the cluster of hits at a low percentile and a high ROC AUC (**Table 1**). Furthermore, these categories can be broken down even more specifically by looking at amino acid identity (e.g., “Alanine,” “Glutamic Acid,” “Tryptophan”). In an example of mutations from Lysine to Leucine (**Fig. 3B**), the reverse is true where the mutations are biased by MM/GBSA. Ultimately, this suggests that the bias needs to be accounted for in each of these methods.

**Figure 3.**
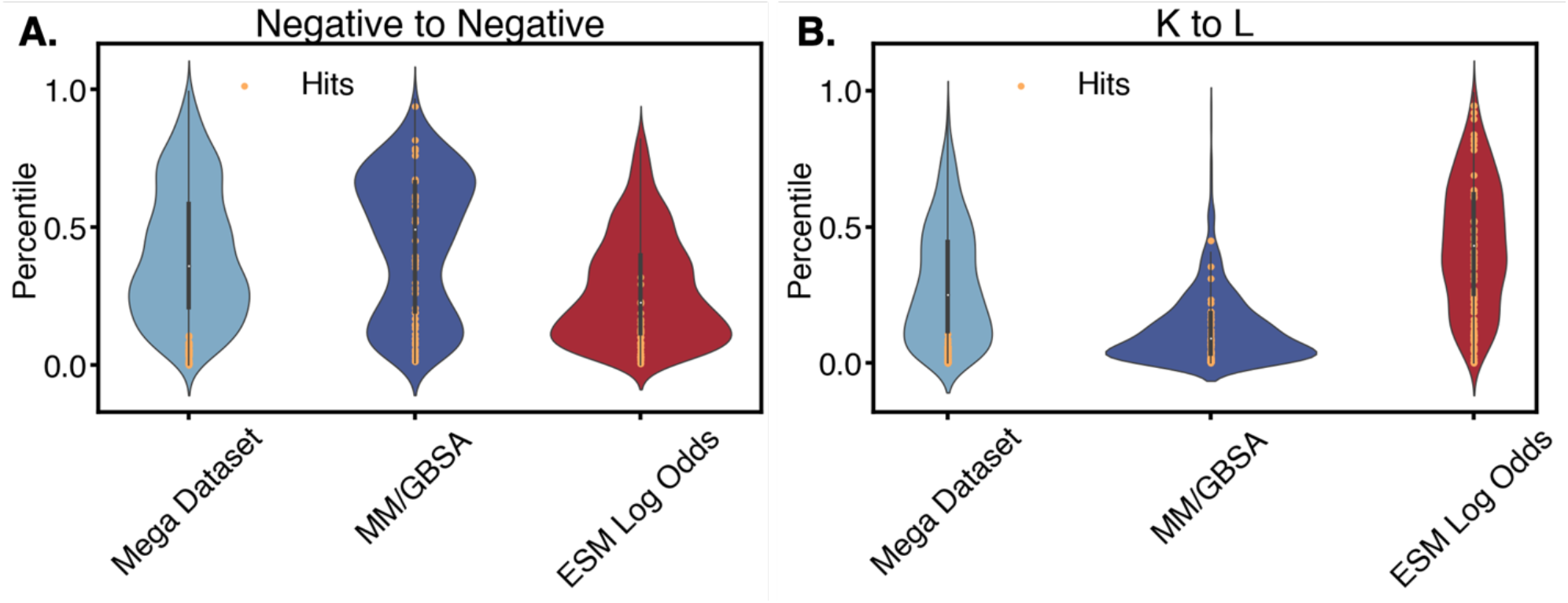
Percentile Distributions by Method. Violin plots depicting the distribution of mutation ranks (percentiles) derived from experiment (values in the Mega Dataset) or predicted by MM/GBSA, and ESM-1b. A) Mutations grouped by physicochemical properties (e.g., “Negative,” “Hydrophobic,” “Non-polar”) and B) mutations grouped by amino acid identity of individual amino acid substitutions (e.g., “Alanine,” “Glutamic Acid,” “Tryptophan”). Hits, based on a 0.5 kcal/mole cut-off, are shown in orange. Graphs depicting all of the groups and amino acid substitutions can be found in **Appendix B**.

### 3.5. Hybrid mutant prioritization

The strong biases exhibited by different methods in several mutation types suggested that a linear combination of percentile outputs (**Eqn. 2**), weighted by mutation type, could optimize the prioritization. This approach generates a new percentile, effectively reordering mutations by integrating information from both methods (**Fig. 4A**).

**Figure 4.**
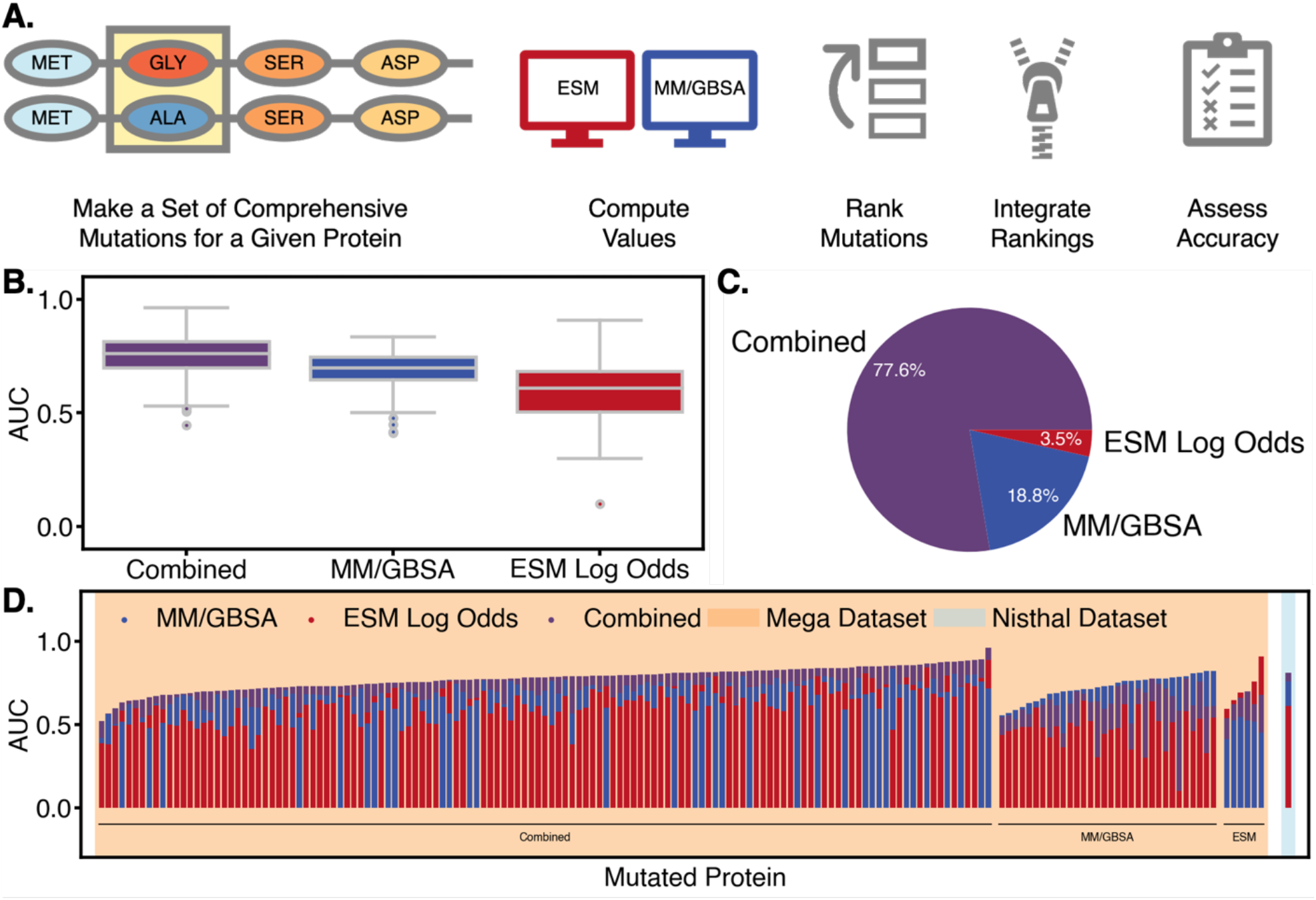
Hybrid mutation prioritization approach assessment. A) Workflow Schematic: the mutation prioritization workflow used to leverage both ESM-1b and MM/GBSA to identify stabilizing mutations. B) Aggregated Performance: a comparison of the performance of the three methods - Combined, MM/GBSA, and ESM Log Odds - using the aggregated area under the curve as the key metric. C) Protein Aggregated Performance: The percentage of all proteins (among both datasets) for which a specific method achieved the highest AUC. D) Detailed AUC Representation: a more granular view of AUC performance in which each protein in the analyzed datasets is represented by three color-coded points along the vertical axis: Blue points show MM/GBSA prediction results, red points indicate ESM log odd predictions and black represent the combined method’s performance. The background color distinguishes the dataset analyzed: pink for the Mega Dataset and light blue for the Nisthal Dataset. For easier visualization, the datapoints are organized into three groups from left to right. The largest group on the left highlights proteins where the Combined method performed best, the next group comprises proteins for which MM/GBSA performed best, and the third group consists of proteins for which ESM Log Odds showed best prediction performance. Of the proteins in the Mega Dataset, ten (PDBs: 2KVS, 2K5H, 1H8K, 6IWS, 1OPS, 3V1A, 2LUM, 2KXD, 1E6H, 2EXD) lacked stabilizing mutations 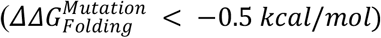 and were consequently excluded from these analyses.

The combined prioritization strategy achieved a higher overall ROC AUC across the dataset than either method alone (**Fig. 4B**). Although individual methods occasionally provided optimal ranking for specific proteins, the combined approach demonstrated superior or comparable performance in most cases (77.6% of the time) (**Fig. 4C**). Notably, the combined method counteracted the biases of both models, and was thus never the worst method for any protein in the dataset and ESM Log Odds was most commonly the worst (76.5% of the time) compared with MM/GBSA (23.5% of the time) (**Fig. 4D**). Additionally, the performance of the combined prioritization strategy was evaluated among the different mutation groups (**Table S4**). Improved ROC AUC values across most mutations groups with the combined prioritization strategy indicate Simpson’s paradox is not being encountered in this analysis.

Due to the speed of obtaining results from PLMs, these findings suggest that integrating complementary insights from physics-based and evolution-based computational tools can enhance the accuracy of mutation prioritization for protein engineering without significantly increasing the computational expense. Additionally, in cases where the PBMs do not contribute significantly to the ordering of mutations, a PLM could replace this tool, significantly accelerating mutation prioritization.

## 4. Discussion and Conclusion

Ultimately, MM/GBSA more closely aligned with the experimental values and trends compared to ESM-1b. Given ESM-1b was not designed to identify stabilizing mutations, its poor performance at this task can be expected. When examining populations of stabilizing mutations and destabilizing mutations, MM/GBSA performed better than ESM-1b. MM/GBSA is sensitive, specific, and accurate, while ESM-1b is less so. MM/GBSA was able to more clearly distinguish *mutable sites* as sites with low standard deviation across the computed computational values, indicative of sites tolerable to a variety of mutations. There are clear quantitative differences between ESM-1b’s and MM/GBSA’s predictions.

Ranking of mutations on a per protein basis was more effective than ranking of mutations on a per position basis for both MM/GBSA and ESM-1b. Therefore, mutational hotspots can be identified both structurally and from sequence data alone. This suggests that both methods are better at identifying mutational hotspots than they are at ranking relative mutations at a particular position, reinforcing the idea that both methods can be utilized as a screening step for higher-accuracy methods.

However, both MM/GBSA and ESM-1b contain bias for particular mutation types. These biases are demonstrated by increased numbers of mutations captured in the top 20% of mutations in a protein and the top 5 mutations at a site, but a relatively similar ROC AUC across mutation types. Limited bias would result in mutation capture being directed correlated with ROC AUC.

The underlying methodology in these approaches can explain the biases in prediction. For both methods, the abstraction of structural information effectively discards biochemically significant context. The implicit solvent model in MM/GBSA cannot account for changes in solvent reorganization and entropy, which can contribute to overestimation and overprediction of mutations residues with charged side chains. Meanwhile, the complete absence of structural information within ESM-1b precludes any accounting for protein fold-induced interatomic interactions. For example, failure to account for the differences between core and surface mutations could result in an overestimation and overprediction of mutations residues with hydrophobic side chains.

When MM/GBSA and ESM-1b results are combined, the overall ROC AUC improves to 0.744 across the whole dataset compared to either MM/GBSA alone (0.684) or ESM Log Odds alone (0.597). The errors of each method are not perfectly correlated so we expect to reduce error, on average, when using an ensemble of the methods. Additionally, the combined method is the best method for prioritization for 77.6% of the proteins in the dataset. A hybrid combined approach consistently outperforms either method alone due to its ability to combine information gleaned from each.

Each computational method for prioritizing stabilizing mutations should be used with a clear awareness of its benefits and drawbacks. MM/GBSA provides consistent values rooted in physics, resulting in a generally applicable method that balances computational cost with accuracy. On the other hand, ESM-1b quickly explores a space with limited understanding of the physical mechanisms, highlighting some mutations that can achieve experimental goals, but cannot extrapolate to all systems. Together, a hybrid approach supplements the physical explanation (MM/GBSA value) for a stabilizing mutation with evolutionary context (ESM log odds value) resulting in both a bolstered understanding of the mutations as well as improved performance. Overall knowledge-based methods such as PLMs can supplement PBMs to better predict experimental measurements.

## References

1. Rigoldi F, Donini S, Redaelli A, Parisini E, Gautieri A. Review: Engineering of thermostable enzymes for industrial applications. APL Bioengineering. 2018;2(1):011501. doi:10.1063/1.4997367

2. McConnell AD, Zhang X, Macomber JL, et al. A general approach to antibody thermostabilization. mAbs. 2014;6(5):1274–1282. doi:10.4161/mabs.29680

3. Xu Z, Cen YK, Zou SP, Xue YP, Zheng YG. Recent advances in the improvement of enzyme thermostability by structure modification. Critical Reviews in Biotechnology. 2020;40(1):83–98. doi:10.1080/07388551.2019.1682963

4. Lutz S, Iamurri SM. Protein Engineering: Past, Present, and Future. In: Bornscheuer UT, Höhne M, eds. Protein Engineering. Vol 1685. Methods in Molecular Biology. Springer New York; 2018:1–12. doi:10.1007/978-1-4939-7366-8_1

5. d’Oelsnitz S, Ellington A. Continuous directed evolution for strain and protein engineering. Current Opinion in Biotechnology. 2018;53:158–163. doi:10.1016/j.copbio.2017.12.020

6. Montanucci L, Capriotti E, Frank Y, Ben-Tal N, Fariselli P. DDGun: an untrained method for the prediction of protein stability changes upon single and multiple point variations. BMC Bioinformatics. 2019;20(Suppl 14):335. doi:10.1186/s12859-019-2923-1

7. Worth CL, Preissner R, Blundell TL. SDM--a server for predicting effects of mutations on protein stability and malfunction. Nucleic Acids Res. 2011;39(Web Server issue):W215–222. doi:10.1093/nar/gkr363

8. Schymkowitz J, Borg J, Stricher F, Nys R, Rousseau F, Serrano L. The FoldX web server: an online force field. Nucleic Acids Res. 2005;33(Web Server issue):W382–388. doi:10.1093/nar/gki387

9. Kellogg EH, Leaver‐Fay A, Baker D. Role of conformational sampling in computing mutation‐induced changes in protein structure and stability. Proteins. 2011;79(3):830–838. doi:10.1002/prot.22921

10. Scarabelli G, Oloo EO, Maier JKX, Rodriguez-Granillo A. Accurate Prediction of Protein Thermodynamic Stability Changes upon Residue Mutation using Free Energy Perturbation. Journal of Molecular Biology. 2022;434(2):167375. doi:10.1016/j.jmb.2021.167375

11. Lihan M, Lupyan D, Oehme D. Target‐template relationships in protein structure prediction and their effect on the accuracy of thermostability calculations. Protein Science. 2023;32(2):e4557. doi:10.1002/pro.4557

12. Li B, Yang YT, Capra JA, Gerstein MB. Predicting changes in protein thermodynamic stability upon point mutation with deep 3D convolutional neural networks. PLoS Comput Biol. 2020;16(11):e1008291. doi:10.1371/journal.pcbi.1008291

13. Benevenuta S, Pancotti C, Fariselli P, Birolo G, Sanavia T. An antisymmetric neural network to predict free energy changes in protein variants. J Phys D: Appl Phys. 2021;54(24):245403. doi:10.1088/1361-6463/abedfb

14. Mansoor S, Baek M, Juergens D, Watson JL, Baker D. Zero‐shot mutation effect prediction on protein stability and function using ROSETTAFOLD. Protein Science. 2023;32(11):e4780. doi:10.1002/pro.4780

15. Pires DEV, Ascher DB, Blundell TL. DUET: a server for predicting effects of mutations on protein stability using an integrated computational approach. Nucleic Acids Res. 2014;42(Web Server issue):W314–319. doi:10.1093/nar/gku411

16. Cheng J, Randall A, Baldi P. Prediction of protein stability changes for single‐site mutations using support vector machines. Proteins. 2006;62(4):1125–1132. doi:10.1002/prot.20810

17. Savojardo C, Fariselli P, Martelli PL, Casadio R. INPS-MD: a web server to predict stability of protein variants from sequence and structure. Bioinformatics. 2016;32(16):2542–2544. doi:10.1093/bioinformatics/btw192

18. Capriotti E, Fariselli P, Casadio R. I-Mutant2.0: predicting stability changes upon mutation from the protein sequence or structure. Nucleic Acids Research. 2005;33(Web Server):W306–W310. doi:10.1093/nar/gki375

19. Rao R, Meier J, Sercu T, Ovchinnikov S, Rives A. Transformer protein language models are unsupervised structure learners. Preprint posted online December 15, 2020. doi:10.1101/2020.12.15.422761

20. Hie BL, Shanker VR, Xu D, et al. Efficient evolution of human antibodies from general protein language models. Nat Biotechnol. 2024;42(2):275–283. doi:10.1038/s41587-023-01763-2

21. Shivakumar D, Williams J, Wu Y, Damm W, Shelley J, Sherman W. Prediction of Absolute Solvation Free Energies using Molecular Dynamics Free Energy Perturbation and the OPLS Force Field. J Chem Theory Comput. 2010;6(5):1509–1519. doi:10.1021/ct900587b

22. Alley EC, Khimulya G, Biswas S, AlQuraishi M, Church GM. Unified rational protein engineering with sequence-based deep representation learning. Nat Methods. 2019;16(12):1315–1322. doi:10.1038/s41592-019-0598-1

23. Rives A, Meier J, Sercu T, et al. Biological structure and function emerge from scaling unsupervised learning to 250 million protein sequences. Proc Natl Acad Sci USA. 2021;118(15):e2016239118. doi:10.1073/pnas.2016239118

24. Heinzinger M, Littmann M, Sillitoe I, Bordin N, Orengo C, Rost B. Contrastive learning on protein embeddings enlightens midnight zone. Preprint posted online November 15, 2021. doi:10.1101/2021.11.14.468528

25. Lin Z, Akin H, Rao R, et al. Evolutionary-scale prediction of atomic-level protein structure with a language model. Science. 2023;379(6637):1123–1130. doi:10.1126/science.ade2574

26. UniProt Consortium. UniParc. https://www.ebi.ac.uk/uniprot/uniparc/

27. Genheden S, Ryde U. The MM/PBSA and MM/GBSA methods to estimate ligand-binding affinities. Expert Opinion on Drug Discovery. 2015;10(5):449–461. doi:10.1517/17460441.2015.1032936

28. Nisthal A, Wang CY, Ary ML, Mayo SL. Protein stability engineering insights revealed by domain-wide comprehensive mutagenesis. Proc Natl Acad Sci USA. 2019;116(33):16367–16377. doi:10.1073/pnas.1903888116

29. Gallagher T, Alexander P, Bryan P, Gilliland GL. Two crystal structures of the B1 immunoglobulin-binding domain of streptococcal protein G and comparison with NMR. Biochemistry. 1994;33(15):4721–4729.

30. Tsuboyama K, Dauparas J, Chen J, et al. Mega-scale experimental analysis of protein folding stability in biology and design. Nature. 2023;620(7973):434–444. doi:10.1038/s41586-023-06328-6

31. Khan S, Vihinen M. Performance of protein stability predictors. Hum Mutat. 2010;31(6):675–684. doi:10.1002/humu.21242

32. Schrödinger Release 2023–4. Published online 2023.

33. Søndergaard CR, Olsson MHM, Rostkowski M, Jensen JH. Improved Treatment of Ligands and Coupling Effects in Empirical Calculation and Rationalization of p K a Values. J Chem Theory Comput. 2011;7(7):2284–2295. doi:10.1021/ct200133y

34. Olsson MHM, Søndergaard CR, Rostkowski M, Jensen JH. PROPKA3: Consistent Treatment of Internal and Surface Residues in Empirical p K a Predictions. J Chem Theory Comput. 2011;7(2):525–537. doi:10.1021/ct100578z

35. Lu C, Wu C, Ghoreishi D, et al. OPLS4: Improving Force Field Accuracy on Challenging Regimes of Chemical Space. J Chem Theory Comput. 2021;17(7):4291–4300. doi:10.1021/acs.jctc.1c00302

36. Beard H, Cholleti A, Pearlman D, Sherman W, Loving KA. Applying Physics-Based Scoring to Calculate Free Energies of Binding for Single Amino Acid Mutations in Protein-Protein Complexes. Salsbury, Jr F, ed. PLoS ONE. 2013;8(12):e82849. doi:10.1371/journal.pone.0082849

