## Supplementary Information for "Prioritizing Stability-enhancing Mutations using a Protein Language Model in conjunction with Physics-Based Predictions"

### 6. Appendix A. Supplementary Information

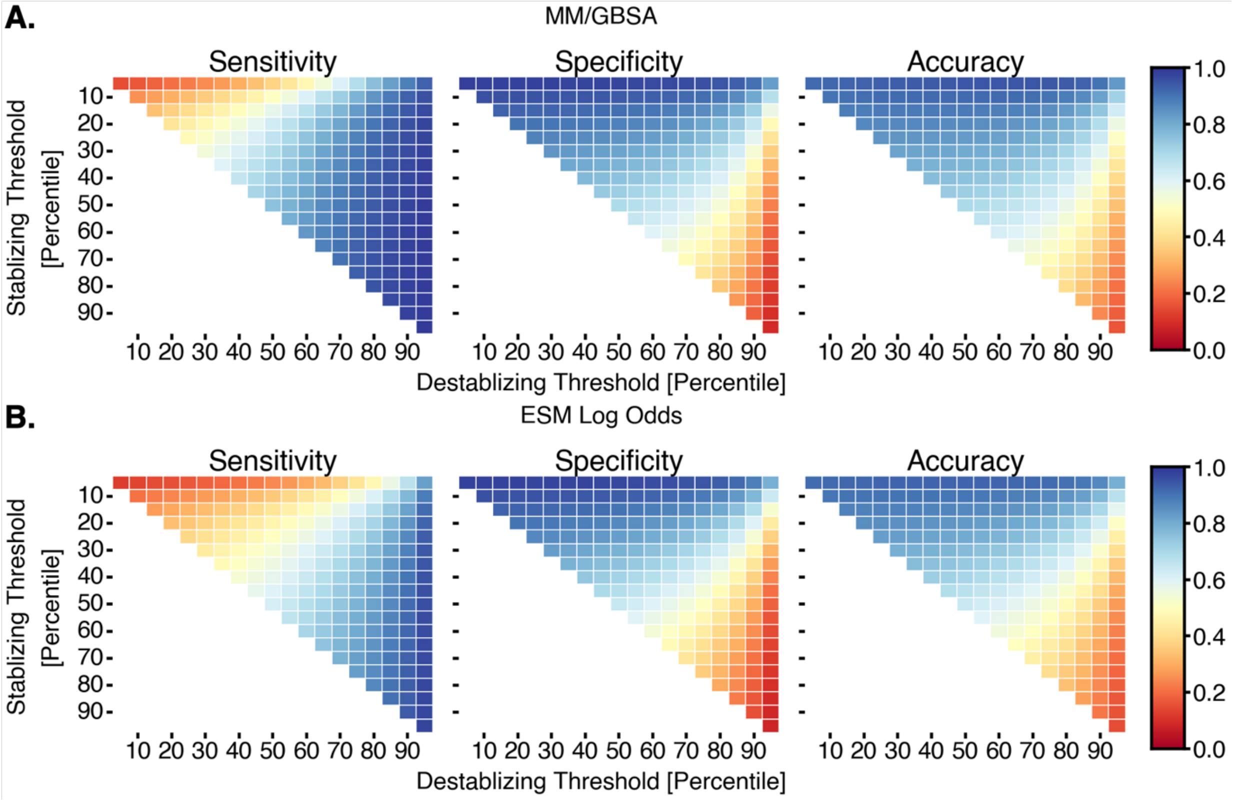


**Figure S1. Sensitivity, Specificity and Accuracy Analysis**. The sensitivity, specificity and accuracy values were graphed while scanning across various percentile thresholds for both the stabilizing and destabilizing categorization. A) For the MM/GBSA values, the top 5% of mutations being stabilizing and bottom 10% of mutations being destabilizing resulted in the highest sensitivity-specificity combination, while for ESM Log Odds, the top 5% of mutations being stabilizing and bottom 5% of mutations being destabilizing resulted in the highest sensitivity-specificity combination. From a conservative perspective the top 5% of mutations being stabilizing and bottom 5% of mutations being destabilizing was taken to be the rule for computationally determining mutation categorization.

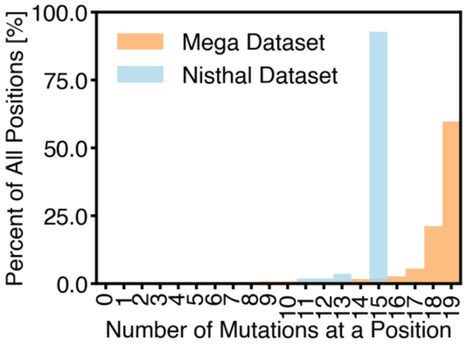


**Figure S2. Number of Mutations per Position in Each Dataset**. The number of mutation values at a position in either the Mega Dataset or the Nisthal Dataset. The maximum number of mutations for the Mega Dataset is 19 mutations, while the maximum number of mutations is 15 for the Nisthal Dataset.

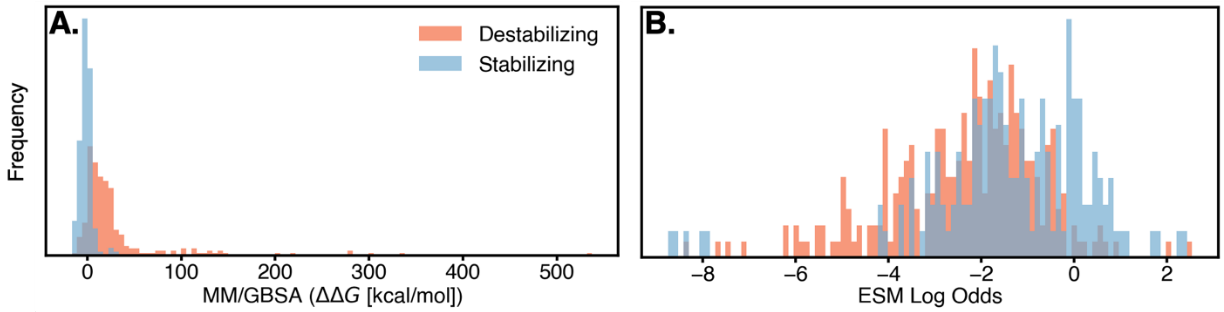


C.

| | MM/GBSA | | ESM Log Odds | |
|---|---|---|---|---|
| | KS Test Statistic | KS Test p value | KS Test Statistic | KS Test p value |
| Stabilizing vs Destabilizing – Mega Dataset | 0.454 | 0.000 | 0.229 | 2.12E-314 |
| Stabilizing vs Destabilizing – Nisthal Dataset | 0.709 | 3.87E-49 | 0.255 | 3.32E-06 |

**Figure S3. Distinguishing stabilizing mutations**. The population of mutation values measured by A) MM/GBSA, and B) ESM log-odds ratio for stabilizing (blue) and destabilizing (orange) mutations, in the B1 immunoglobulin-binding domain of streptococcal protein G. C) Significant differences in populations are indicated by the p-value of a KS test and the degree of separation by the KS statistic.

A.

| | MM/GBSA | | ESM Log Odds | |
|---|---|---|---|---|
| | Spearman | Pearson | Spearman | Pearson |
| Rank all Mutations for the Whole Protein - Mega Dataset | 0.563 | 0.597 | 0.436 | 0.459 |
| Rank all Mutations for the Whole Protein – Nisthal Dataset | 0.635 | 0.635 | 0.237 | 0.237 |
| Rank the Mutations at each Position - Mega Dataset | 0.323 | 0.323 | 0.239 | 0.239 |
| Rank the Mutations at each Position – Nisthal Dataset | 0.430 | 0.430 | 0.125 | 0.125 |

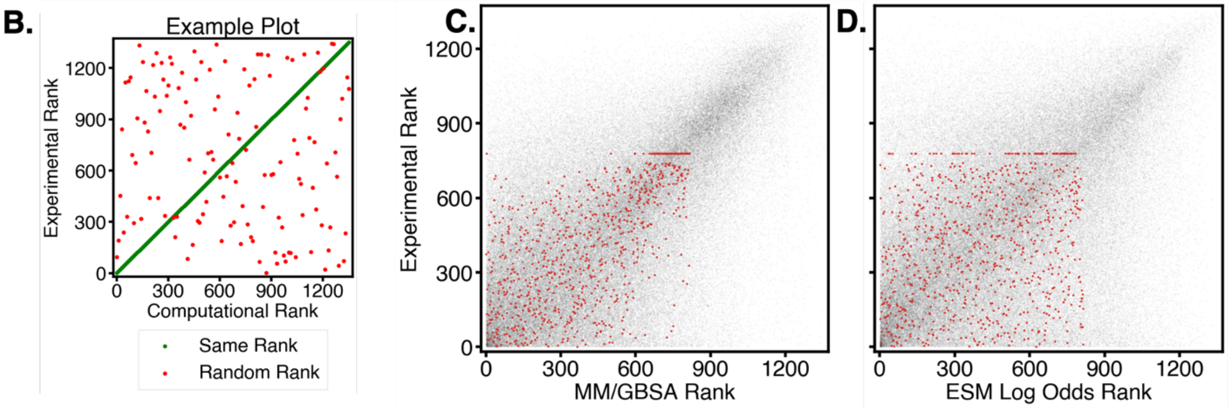


**Figure S4. Mutation ranking**. A) Spearman and Pearson correlations of the ranking performed by each computational method compared to the ranking from experimental values, where the ranking was either for the entire protein or on a per-position basis. B) Visual representation indicating a matching vs. random ranking between experimental and computational values. C-D) The experimental mutational rank versus the C) MM/GBSA, and D) ESM log-odds ratios mutational rankings for either the B1 immunoglobulin-binding domain of streptococcal protein G (red) or the entire Mega Dataset (black) on a per protein basis.

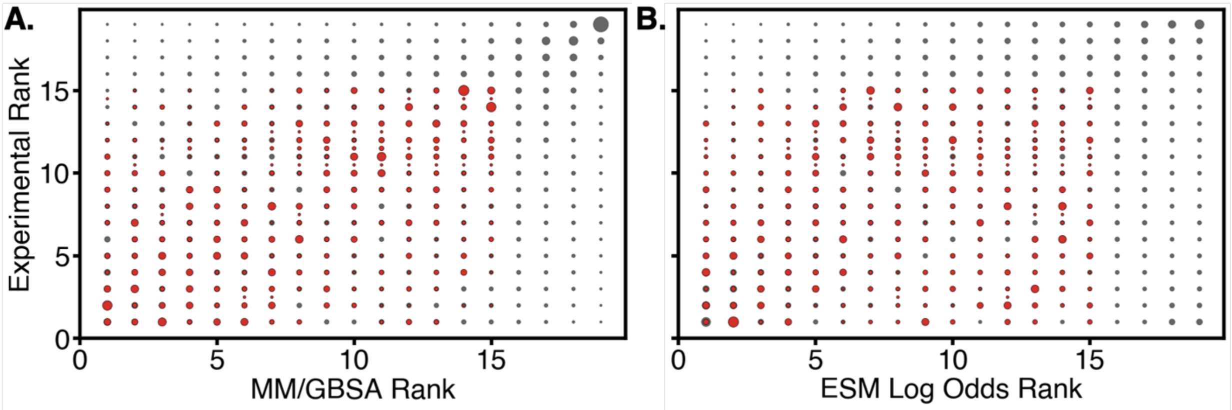


**Figure S5. Mutation ranking per position**. The experimental rank versus the A) MM/GBSA, and B) ESM log-odds ratios position rankings for either the B1 immunoglobulin-binding domain of streptococcal protein G (red) or the entire Mega Dataset (black), where the size of the dot indicates the number of points with those two rankings.

**Table S1. Mega Dataset truth table**.

| | MM/GBSA | ESM Log Odds |
|---|---|---|
| True Positives (Exp. < -0.5 kcal/mol, Comp. Beneficial) | 1081 | 892 |
| True Negatives (Exp. > 0.5 kcal/mol, Comp. Detrimental) | 7779 | 6742 |
| False Positives (Exp. > 0.5 kcal/mol, Comp. Beneficial) | 1620 | 1738 |
| False Negatives (Exp. < -0.5 kcal/mol, Comp. Detrimental) | 85 | 197 |
| True Neutral (Exp. > -0.5 kcal/mol & < 0.5 kcal/mol, Comp. Neutral) | 73691 | 72659 |
| False Neutral Positive (Exp. < -0.5 kcal/mol, Comp. Neutral) | 6170 | 6258 |
| False Neutral Negative (Exp. > 0.5 kcal/mol, Comp. Neutral) | 76568 | 77512 |
| False Positive Neutral (Exp. > -0.5 kcal/mol & < 0.5 kcal/mol, Comp. Beneficial) | 5821 | 5892 |
| False Negative Neutral (Exp. > -0.5 kcal/mol & < 0.5 kcal/mol, Comp. Detrimental) | 837 | 1762 |
| Sensitivity | 0.927 | 0.819 |
| Specificity | 0.828 | 0.795 |
| Accuracy | 0.839 | 0.798 |

**Table S2. B1 immunoglobulin-binding domain of streptococcal protein G truth table.**

| | MM/GBSA | ESM Log Odds |
|---|---|---|
| True Positives (Exp. < -0.5 kcal/mol, Comp. Beneficial) | 18 | 14 |
| True Negatives (Exp. > 0.5 kcal/mol, Comp. Detrimental) | 35 | 21 |
| False Positives (Exp. > 0.5 kcal/mol, Comp. Beneficial) | 4 | 5 |
| False Negatives (Exp. < -0.5 kcal/mol, Comp. Detrimental) | 0 | 5 |
| True Neutral (Exp. > -0.5 kcal/mol & < 0.5 kcal/mol, Comp. Neutral) | 364 | 352 |
| False Neutral Positive (Exp. < -0.5 kcal/mol, Comp. Neutral) | 143 | 142 |
| False Neutral Negative (Exp. > 0.5 kcal/mol, Comp. Neutral) | 226 | 239 |
| False Positive Neutral (Exp. > -0.5 kcal/mol & < 0.5 kcal/mol, Comp. Beneficial) | 18 | 21 |
| False Negative Neutral (Exp. > -0.5 kcal/mol & < 0.5 kcal/mol, Comp. Detrimental) | 6 | 15 |
| Sensitivity | 1.000 | 0.737 |
| Specificity | 0.897 | 0.808 |
| Accuracy | 0.930 | 0.778 |

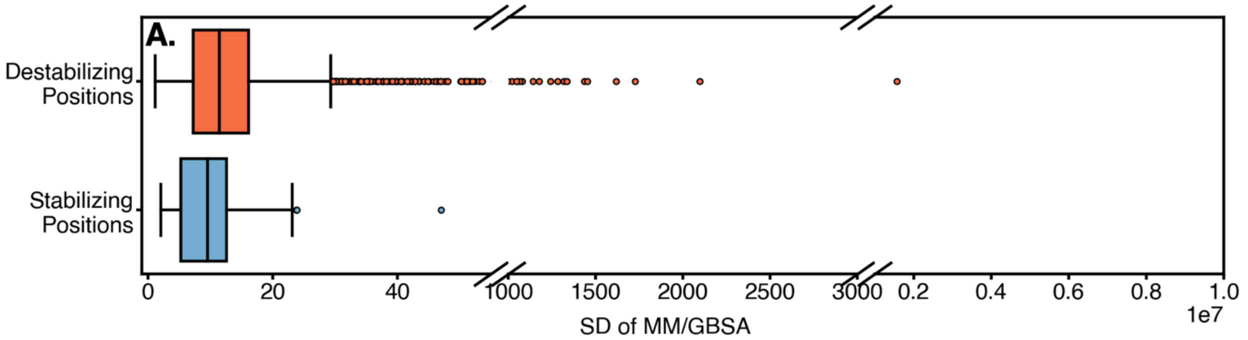


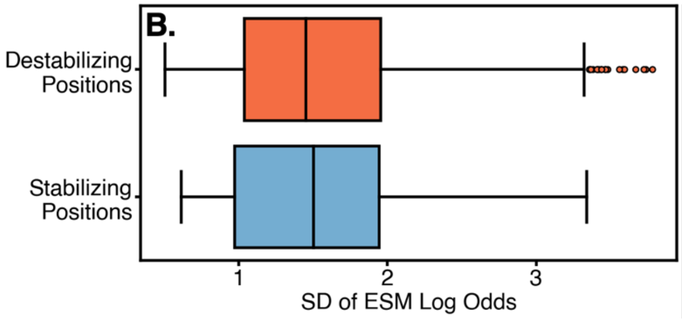


C.

| | MM/GBSA | ESM Log Odds |
|---|---|---|
| KS Test Statistic | 0.172 | 0.0725 |
| KS Test p Value | 0.00195 | 0.555 |

**Figure S6. Distinguishing *mutable sites***. The standard deviation of mutation values measured by A) MM/GBSA, and B) ESM log-odds ratios for the entire Mega Dataset (N = 179 proteins). C) The KS test statistics and p-values between each of the populations (destabilizing and stabilizing mutations) for each computational technique.

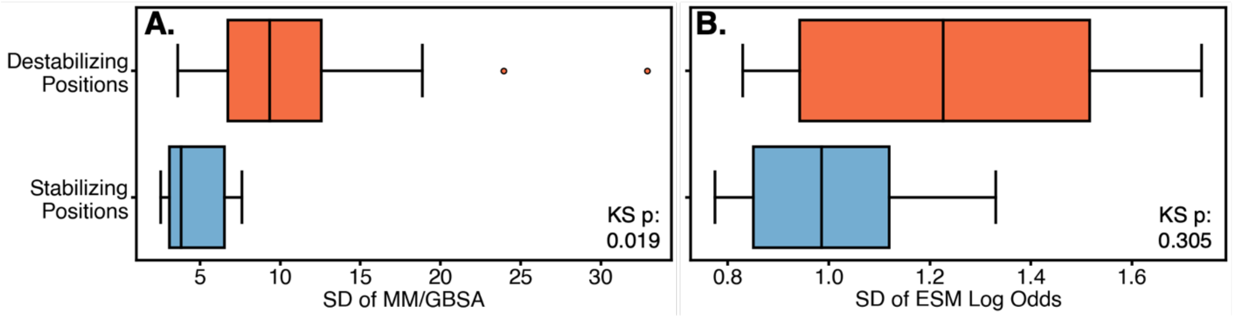


**Figure S7. Distinguishing *mutable sites***. The standard deviation (SD) of mutation-induced free energy/score changes computed via A) MM/GBSA, and B) ESM log-odds ratio for mutated positions in the Nisthal Dataset. Significant differences in populations are indicated by the p-value of a KS test.

**Table S3. Ranking of MM/GBSA- and ESM-1b-captured stabilizing mutations by categories of amino acid changes for the Nisthal Dataset**.

| | Exp. Hit | MM/GBSA | | | ESM Log Odds | | |
|---|---|---|---|---|---|---|---|
| | | Top 20% (Protein) | Top 5 (Position) | ROC AUC | Top 20% (Protein) | Top 5 (Position) | ROC AUC |
| Positive to Negative | 2 | 0.00% | 0.00% | 1.00 | 50.0% | 100% | 0.700 |
| Negative to Positive | 6 | 50.0% | 33.3% | 0.681 | 33.3% | 50.0% | 0.764 |
| Positive to Non-charged Polar | 7 | 42.9% | 42.9% | 0.832 | 14.3% | 42.9% | 0.460 |
| Negative to Non-charged Polar | 16 | 37.5% | 12.5% | 0.579 | 25.0% | 25.0% | 0.581 |
| Non-charged Polar to Positive | 6 | 33.3% | 50.0% | 0.500 | 33.3% | 50.0% | 0.706 |
| Non-charged Polar to Negative | 2 | 0.00% | 0.00% | 0.868 | 100.0% | 100.0% | 0.941 |
| Negative to Hydrophobic | 19 | 26.3% | 63.2% | 0.526 | 10.5% | 15.8% | 0.576 |
| Hydrophobic to Negative | 4 | 0.0% | 25.0% | 0.895 | 75.0% | 50.0% | 0.697 |
| Positive to Hydrophobic | 10 | 60.0% | 70.0% | 0.781 | 20.0% | 10.0% | 0.484 |
| Hydrophobic to Positive | 10 | 30.0% | 50.0% | 0.794 | 20.0% | 30.0% | 0.539 |
| Non-charged Polar to Hydrophobic | 29 | 51.7% | 55.2% | 0.724 | 44.8% | 48.3% | 0.703 |
| Hydrophobic to Non-charged Polar | 9 | 22.2% | 22.2% | 0.823 | 33.3% | 33.3% | 0.562 |
| Non-charged Polar to Non-charged Polar | 9 | 66.7% | 33.3% | 0.767 | 22.2% | 22.2% | 0.820 |
| Hydrophobic to Hydrophobic | 24 | 37.5% | 58.3% | 0.865 | 37.5% | 37.5% | 0.552 |
| Negative to Negative | 4 | 75.0% | 25.0% | 0.750 | 100.0% | 100.0% | 0.958 |
| Positive to Positive | 4 | 100% | 100% | 0.500 | 0.00% | 0.00% | 0.125 |
| Any to Proline | 0 | | | | | | |
| Proline to Any | 0 | | | | | | |
| Any to Cysteine | 0 | | | | | | |
| Cysteine to Any | 0 | | | | | | |
| Small to Large | 29 | 44.8% | 37.9% | 0.710 | 37.9% | 37.9% | 0.715 |
| Large to Small | 6 | 0.00% | 33.3% | 0.848 | 100% | 83.3% | 0.854 |
| Any to Any | 161 | 41.6% | 46.6% | 0.758 | 32.3% | 36.0% | 0.609 |

Each row is a different category of mutation where the first word describes the wildtype residue which is then mutated to another amino acid described by the second word (for instance, "positive to hydrophobic" indicates the wildtype amino acid is positive and it is then mutated to a hydrophobic residue). The columns indicate the percent of hits the computational method captured in the "Top 20% (Protein") and "Top 5 (Position)" as well as the "ROC AUC" for all mutations of a given category.

**Table S4. Ranking of the combined prioritization strategy captured stabilizing mutations by categories of amino acid changes for the Mega Dataset**.

| | Exp. Hit | Combined | | | MM/GBSA | | | ESM Log Odds | | |
|---|---|---|---|---|---|---|---|---|---|---|
| | | Top 20% (Protein) | Top 5 (Position) | ROC AUC | Top 20% (Protein) | Top 5 (Position) | ROC AUC | Top 20% (Protein) | Top 5 (Position) | ROC AUC |
| Positive to Negative | 113 | 31.9% | 7.96% | 0.70 | + | + | + | + | + | - |
| Negative to Positive | 237 | 51.5% | 29.1% | 0.73 | - | + | - | - | + | - |
| Positive to Non-charged Polar | 248 | 29.8% | 12.5% | 0.65 | + | + | - | - | + | - |
| Negative to Non-charged Polar | 468 | 46.6% | 20.1% | 0.68 | + | - | - | - | + | - |
| Non-charged Polar to Positive | 238 | 56.3% | 26.9% | 0.71 | - | + | - | + | + | - |
| Non-charged Polar to Negative | 150 | 46.7% | 28.7% | 0.70 | - | - | - | + | + | - |
| Negative to Hydrophobic | 896 | 47.7% | 23.9% | 0.69 | + | + | - | - | - | - |
| Hydrophobic to Negative | 148 | 33.8% | 17.6% | 0.82 | - | - | - | + | + | - |
| Positive to Hydrophobic | 639 | 52.0% | 29.7% | 0.67 | + | - | - | - | - | - |
| Hydrophobic to Positive | 139 | 32.4% | 23.0% | 0.76 | + | + | - | + | + | - |
| Non-charged Polar to Hydrophobic | 1130 | 60.0% | 34.5% | 0.74 | - | = | - | - | - | - |
| Hydrophobic to Non-charged Polar | 346 | 35.0% | 18.2% | 0.77 | - | - | - | + | + | - |
| Non-charged Polar to Non-charged Polar | 406 | 52.2% | 25.6% | 0.71 | - | - | - | - | + | - |
| Hydrophobic to Hydrophobic | 973 | 41.2% | 31.6% | 0.76 | - | - | - | + | - | - |
| Negative to Negative | 44 | 70.5% | 40.9% | 0.75 | - | - | - | + | + | + |
| Positive to Positive | 31 | 93.5% | 22.6% | 0.65 | - | - | + | - | = | - |
| Any to Proline | 131 | 37.4% | 26.0% | 0.86 | - | - | - | + | + | - |
| Proline to Any | 224 | 26.8% | 25.9% | 0.74 | + | - | - | + | - | - |
| Any to Cysteine | 821 | 63.1% | 56.4% | 0.70 | - | - | - | - | - | - |
| Cysteine to Any | 1 | 100% | 100% | 1.00 | - | - | - | = | - | - |
| Small to Large | 927 | 40.8% | 21.4% | 0.72 | + | + | - | - | - | - |
| Large to Small | 861 | 50.9% | 36.8% | 0.73 | - | - | - | - | - | - |
| Any to Any | 7360 | 48.9% | 29.9% | 0.75 | - | - | - | - | - | - |

Each row is a different category of mutation where the first word describes the wildtype residue which is then mutated to another amino acid described by the second word (for instance, "positive to hydrophobic" indicates the wildtype amino acid is positive and it is then mutated to a hydrophobic residue). The columns indicate the percent of hits the computational method captured in the "Top 20% (Protein") and "Top 5 (Position)" as well as the "ROC AUC" for all mutations of a given category. A comparison between the combined prioritization strategy and MM/GBSA and ESM Log Odds results are indicated in the MM/GBSA and ESM Log Odds columns respectively. A "+" indicates that the individual method has a larger value in this row, while a "-" indicates that the individual method has a smaller value in this row, and a "=" indicates that the individual method has an equal value in this row when MM/GBSA and ESM Log Odds values are compared to the combined prioritization strategy values.

### 7. Appendix B. Supplementary Folder
